# Genomic sequencing identifies secondary findings in a cohort of parent study participants

**DOI:** 10.1101/183186

**Authors:** Michelle L. Thompson, Candice R. Finnila, Kevin M. Bowling, Kyle B. Brothers, Matthew B. Neu, Michelle D. Amaral, Susan M. Hiatt, Kelly M. East, David E. Gray, James M. J. Lawlor, Whitley V. Kelley, Edward J. Lose, Carla A. Rich, Shirley Simmons, Shawn E. Levy, Richard M. Myers, Gregory S. Barsh, E. Martina Bebin, Gregory M. Cooper

**Affiliations:** HudsonAlpha Institute for Biotechnology, Huntsville, AL, USA; University of Louisville, Louisville, KY, USA; University of Alabama at Birmingham, Birmingham, AL, USA

**Keywords:** Secondary findings, genomic sequencing, disease risk, CSER, ACMG

## Abstract

**PURPOSE:** Clinically relevant secondary variants were identified in parents enrolled with a child with developmental delay and intellectual disability.

**METHODS:** Exome/genome sequencing and analysis of 789 ‘unaffected’ parents was performed.

**RESULTS:** Pathogenic/likely pathogenic variants were identified in 21 genes within 25 individuals (3.2%), with 11 (1.4%) participants harboring variation in a gene defined as clinically actionable by the ACMG. Of the 25 individuals, five carried a variant consistent with a previous clinical diagnosis, thirteen were not previously diagnosed but had symptoms or family history with probable association with the detected variant, and seven reported no symptoms or family history of disease. A limited carrier screen was performed yielding 15 variants in 48 (6.1%) parents. Parents were also analyzed as mate-pairs to identify cases in which both parents were carriers for the same recessive disease; this led to one finding in *ATP7B.* Four participants had two findings (one carrier and one non-carrier variant). In total, 71 of the 789 enrolled parents (9.0%) received secondary findings.

**CONCLUSION:** We provide an overview of the rates and types of clinically relevant secondary findings, which may be useful in the design, and implementation of research and clinical sequencing efforts to identify such findings.

## INTRODUCTION

Whole exome and genome sequencing (WES/WGS) have proven to be powerful tests for identifying clinically relevant genetic variation. The existence of secondary and incidental findings has catalyzed debate regarding the types of findings that should be sought by sequencing labs, the circumstances in which certain types of variants should be returned, and the necessary extent of patient consent, education, and genetic counseling. The American College of Medical Genetics and Genomics (ACMG) released recommendations about the interpretation of variants in genes considered to be clinically actionable, including those that pose high risk of cancer and heart disease. The ACMG suggests that these be sought and provided to patients that consent to receive such results ^1, 2^. Recommendations related to use of specific gene lists and approaches for returning secondary findings were intended to be used in clinical contexts, although it is also important to examine them in translational research contexts.

Through a study that was part of the Clinical Sequencing Exploratory Research (CSER) Consortium ^3^, we assessed the utility of WES/WGS to identify genetic causes of developmental delay, intellectual disability (DD/ID), and related congenital anomalies. We have sequenced affected probands from 455 families, and have identified DD/ID-related pathogenic/likely pathogenic (P/LP) variants in 29% of cases ^4^. As our DD/ID study includes proband-parent trios, we have the ability to assess secondary findings in a sizable cohort of adults ^4^.

We use the term ‘secondary findings’ throughout the manuscript to describe variation identified via proactive searching ^5^ and report rates and types of secondary findings in context of reported symptoms or family history. Our experiences and data suggest the value of genomic sequencing in a clinical setting not only for disease patients, but also for those not currently exhibiting an overt disease phenotype. We demonstrate the utility of dissemination of such findings in a cohort of parent study participants, and highlight this through case study analyses.

## METHODS

### Study participant population

There was no public recruitment for this study. Parent and children (n=455 families) participants were enrolled at North Alabama Children’s Specialists in Huntsville, AL. Consent was obtained for study participation and publication of data generated by this study. Review boards at Western Institutional Review Board (20130675) and the University of Alabama at Birmingham (X130201001) approved and monitored this study.

### Patient preferences and consent

We developed the Preferences Instrument for Genomic Secondary Results (PIGSR) ^6^ to elicit parents’ preferences for receiving categories of secondary results. This instrument divides secondary findings into 13 distinct disease categories (Figure 1). Results were returned to parent participants only when they opted to receive secondary findings. Decisions regard disclosure of secondary findings solely in the proband were based on a combination of parent preferences for themselves and medical relevance to the proband during childhood. In the case of adopted probands, preferences were solicited from the adoptive parents on behalf of the proband.

**Figure 1.**
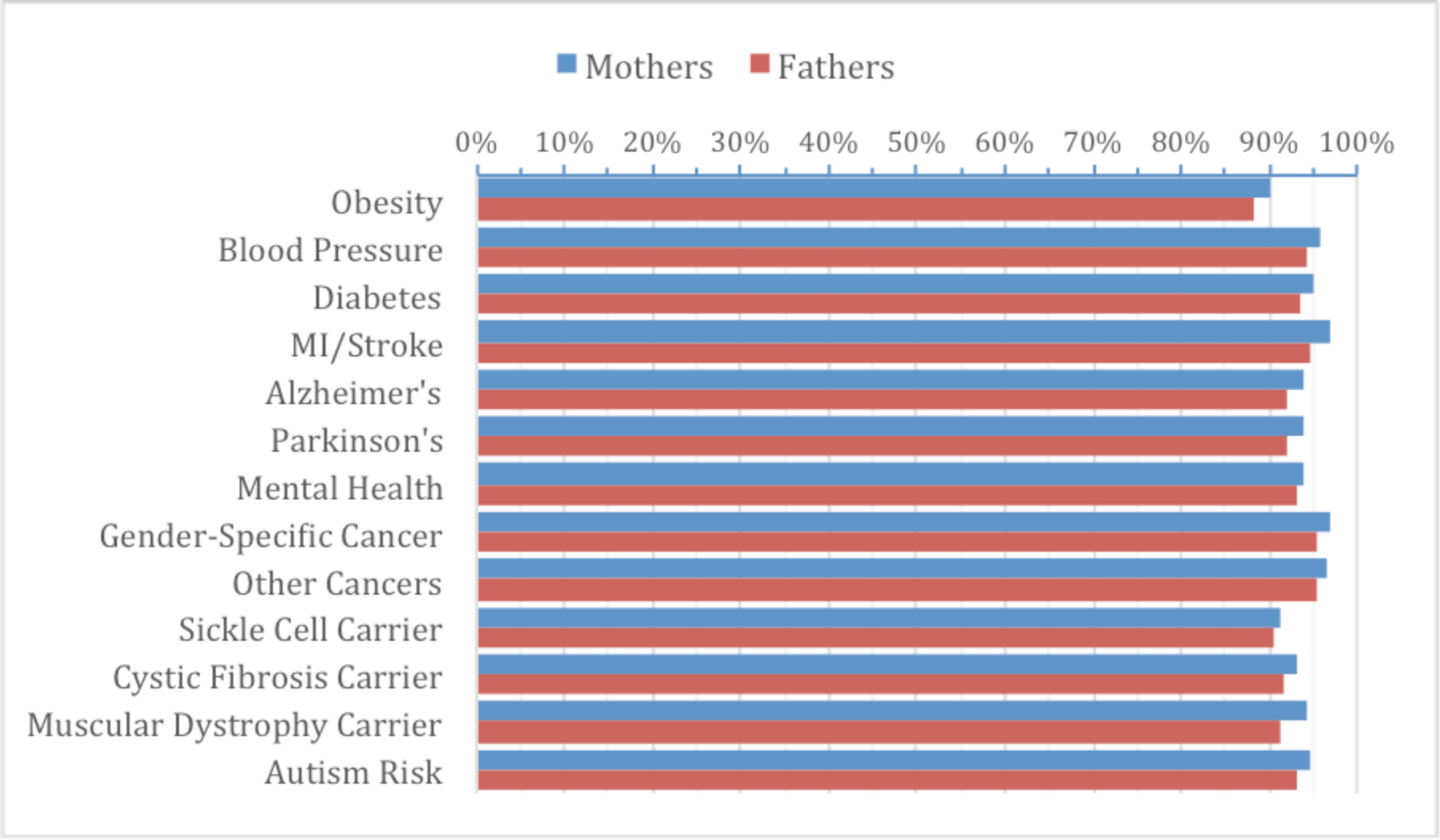
Participant preferences for receipt of secondary genetic findings. Participant preferences were assessed for return of genetic variation across a number of different disease categories. An overwhelmingly large majority (85%) of study participants chose to receive any identified secondary variant, regardless of disease association (n=789).

### Phenotyping

At enrollment, a genetic counselor generated a three-generation pedigree based on family history reported by the parents/guardians of the proband. Parents’ health records were not available to the study nor was a physical exam performed. The genetic counselor asked questions related to family history of cancer and sudden/unusual deaths of adults (e.g. cardiac arrest or aneurysm). Cascade sequencing was not conducted as part of this study. We have (1) retained the language used by the participant and (2) included any reported family history that is plausibly related to the phenotype of concern.

### Return of results

Parent participants that received secondary findings were scheduled for private disclosure with a medical geneticist and genetic counselor. The clinical significance of findings was addressed and documents detailing variant information and relevant resources were provided. Secondary findings were not by default placed in the participant’s medical record and no formal referrals to relevant specialists were made. If the participants chose to share results with their healthcare provider, formal referrals could be coordinated.

### Sequencing and variant information

Further details regarding WES/WGS, read alignment, variant calling, filtering, classification, and validation can be found in our previous report ^4^ and in Supplemental Methods. Briefly, we identified secondary variation in ACMG genes^1, 2^; P/LP variation in ClinVar (not in ACMG genes); recessive variation in individuals who harbored two or more P/LP variants in the same gene; variation in OMIM genes in which both parents of parental pair harbored P/LP variation for the same recessive disorder; and carrier status information in *CFTR, HEXA,* and *HBB.* Only P/LP variants were returned.

### Data sharing

Identified variants in parent participants have been shared through ClinVar and dbGaP, with consent. Additional information is provided in Supplemental Methods.

## RESULTS

### Demographics of study population

Of 455 enrolled families, 424 included at least one parent, and both parents were available for 365 families. Demographics for the 789 parent participants are reported in Table 1. The study population had a mean age of 41 years and included 422 females and 367 males. 80.5% selfreported to be of European ancestry (“White”), 8.5% as African-American (“Black”), and 8.2% as “Other or Multiracial”. Over 25% had a high school diploma or less, while 34.5% reported some college education (Table 1).

**Table 1.**
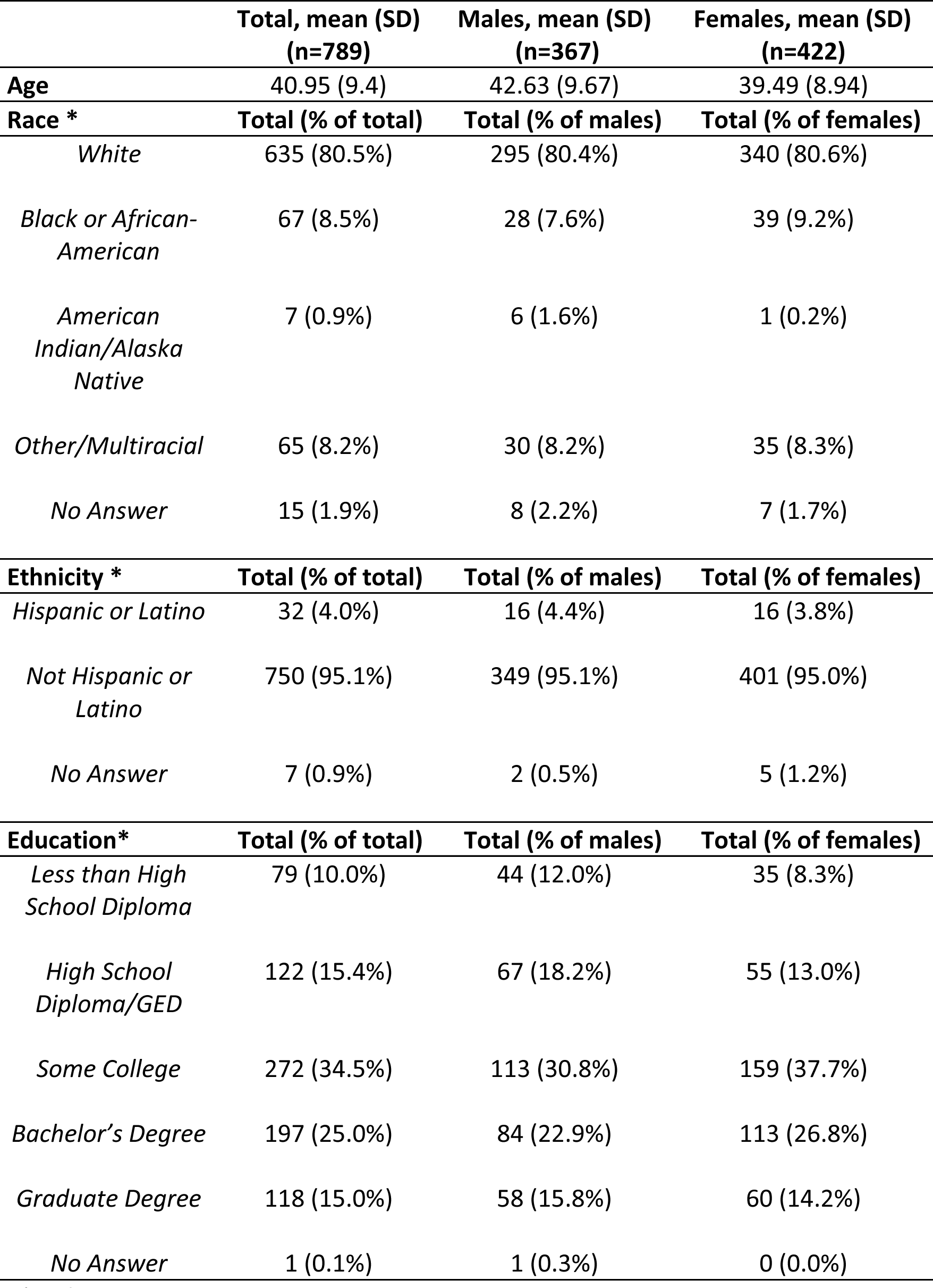
Demographics of parent participants enrolled in the HudsonAlpha CSER project.

*Self-reported

### Patient Preferences

One goal of our study was to understand patient preferences as they relate to receipt of secondary findings across various disease categories ^6^. 85% of parents requested all secondary findings, while 1.6% declined to receive all findings. The most frequently requested category was risk for gender-specific cancers (breast, ovarian, testicular and prostate; n=584, 96.1%). The least frequently requested result was risk for developing obesity (n=542, 89.2%) (Figure 1).

### Carrier status findings

We conducted a limited carrier screen for variants relevant to cystic fibrosis (*CFTR,* MIM: 219700), beta-thalassemia (HBB, MIM: 613985), sickle cell disease (HBB, MIM: 603903), and Tay-Sachs disease (*HEXA,* MIM: 272800), which are among the most common Mendelian diseases (average carrier risk is 1/40) ^7-9^. We observed eight P/LP variants in *CFTR* across 35 individuals (4.4% of parent cohort), four *HEXA* variants across five individuals (0.6%), and three *HBB* variants across eight individuals (1%) (Table 2; Table S2). Additionally, we searched for cases in which parental “mate pairs” were both carriers for variants in a gene associated with a recessive disorder. This analysis led to returnable findings for one of the 365 parental pairs; both parents harbored variation in *ATP7B* associated with Wilson disease (MIM: 277900) (Table S2).

**Table 2.**
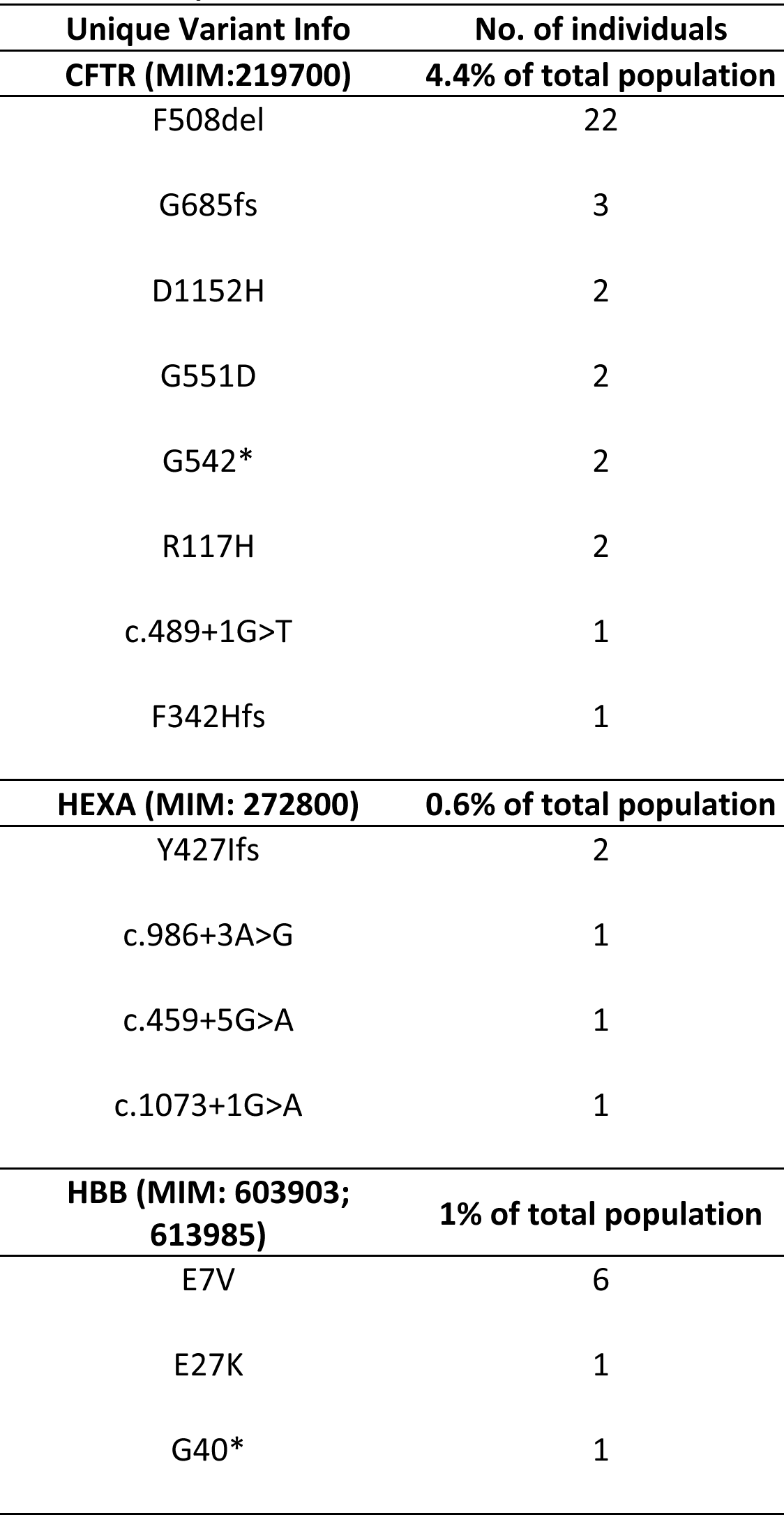
Unique variants of carrier status in CFTR, HEXA, and HBB

### Secondary variation in individuals with a previous clinical diagnosis

P/LP variants were found in five individuals with a self-reported previous clinical diagnosis but in whom a specific genetic cause was unknown. A 35-year-old female individual was found to harbor a heterozygous missense variant in *SLC4A1* (spherocytosis, MIM: 612653), and had family history of related disease (Table 3; Table S1). We identified three missense variants (two likely in *cis*) in *SLC22A5* in a 37-year-old female with recessive systemic primary carnitine deficiency (MIM: 212140). Finally, a canonical splice donor site (D1) variant affecting *PKD2* was identified in a 36-year-old female with polycystic kidney disease (MIM: 613095). This individual also reported a family history of disease (Table 3; Table S1).

**Table 3.** Secondary findings of enrolled parents segregated into “Clinically diagnosed”, “Notable family history and/or symptomatic”, and “Currently asymptomatic with no family history”.

| Age (Male/Female) | Gene | Variant Info | Associated Phenotype (MIM) | Phenotypes or family history reported by parent participants* |
| --- | --- | --- | --- | --- |
| **Clinically diagnosed (0.6% total population)** |  |  |  |  |
| 35- F | *SLC4A1* | V488M | Spherocytosis, type 4 (612653) | Clinically diagnosed with spherocytosis; Two daughters and father with spherocytosis |
| 37- F | *SLC22A5* | A142S; T440M, R488H | Carnitine deficiency, systemic primary (212140) | Clinically diagnosed with carnitine deficiency |
| 36- F | *PKD2* | c.1319+1G>A | Polycystic kidney disease 2 (613095) | Clinically diagnosed with polycystic kidney disease (PKD); mother, brother, 2 nieces, maternal aunt, uncle and grandmother with PKD |
| 30- F | *DSG2* | V986fs | Cardiomyopathy, dilated, 1BB; Arrhythmogenic right ventricular dysplasia 10 (612877; 610193) | Clinically diagnosed with postpartum cardiomyopathy; Paternal family history of arrhythmia; paternal uncle with two “heart attacks” prior to age 40 |
| 52- M | *ANK2* | E1458G | Cardiac arrhythmia, ankryin-B-related, Long QT syndrome 4 (600919) | Clinically diagnosed with hypertrophic cardiomyopathy and arrhythmia; father died with ischemic heart disease |
| **Notable family history and/or symptomatic (1.6% of total population)** |  |  |  |  |
| 29- F | *CLCN1* | F413C | Myotonia congenita, dominant (160800) | Leg cramps and restless legs in childhood, still occasionally has cramps; Mother diagnosed with myotonia congenita, 10 years; Maternal grandfather with a muscle biopsy performed in 30s and “stiffness” especially in cold, 30s |
| 35- F | *MFN2* | W740S | Charcot-Marie-Tooth disease, axonal, type 2A2A (609260) | History of muscle wasting in back, lower extremities; brother clinically diagnosed with CMT, 30s; multiple family members affected with “unspecified muscle disorder” |
| 40- M | *BRCA1* | G1756fs | Breast-ovarian cancer, familial 1 (604370) | Mother with breast cancer, 30s |
| 38- F | *BRCA2* | c.8488-1G>A | Breast-ovarian cancer, familial 2 (612555) | Maternal grandfather with bilateral breast cancer, 60s; Paternal grandmother with breast cancer, age unknown |
| 33- F | *BARD1* | E652fs | Breast cancer susceptibility (114480) | Maternal great-grandmother with breast cancer, 50s; Maternal grandmother had bladder, lung, and peritoneal cancer, age unknown |
| 43- M | *PMS2* | P246fs | Hereditary nonpolyposis colorectal cancer, type 4 (614337) | Father (60s) and paternal aunt (40s) had colon cancer; Paternal aunt (60s) and grandmother (50s) with breast cancer |
| 28- F | *SCN4A* | T1313M | Paramyotonia congenita (168300) | At enrollment, no report of neuromuscular phenotypes. At return of results, indicated that she had muscle stiffness but always thought she was “easily fatigued” and had “low stamina”; Mother displays similar symptoms |
| 41- M | *HARS* | R137Q | Charcot-Marie-Tooth, axonal, type 2W (616625) | At enrollment, no report of neuromuscular phenotypes. At return of results, indicated that he had CMT-associated phenotypes. Always thought he was “just clumsy” |
| 32- F | *KCNQ1* | R366W | Long QT syndrome 1 (192500) | Father with coronary artery disease with triple by-pass, early 50s, paternal aunt with early-onset stroke, late 30s |
| 47- M | *KCNQ1* | P7S | Long QT syndrome 1 (192500)) | Mother “fainted” and “hit the floor”-was told this impact prevented cardiac arrest |
| 39- M | *MYBPC3* | E542Q | Hypertrophic cardiomyopathy 4; Dilated cardiomyopathy 1MM (115197; 615396) | “Leaky heart valve”; Dad has pace maker and mom has “leaky heart valve”, 60s |
| 30- M | *DDX41* | D140fs | Susceptibility to familial myeloproliferative/ lymphoproliferative neoplasms (616871) | Paternal cousin with lymphoma “unspecified”, age unknown |
| 37- F | *MC4R* | C271Y | Obesity, autosomal dominant (601665) | Obese (BMI: 41) |
| **Currently asymptomatic with no family history (0.9% of total population)** |  |  |  |  |
| 52- F | *SCN5A* | T1303M | Long QT syndrome-3 (603830) | Recommended to have cardiovascular evaluation |
| 50-M | *DSG2* | E1020fs | Cardiomyopathy, dilated, 1BB; Arrhythmogenic right ventricular dysplasia 10 (612877; 610193) | Recommended to have cardiovascular evaluation |
| 31-M | *ACTN1* | V105I | Bleeding disorder, platelet type, 15 (615193) | Recommended to have a complete blood count and functional platelet study |
| 33- M | *MSH2* | Y570fs | Hereditary nonpolyposis colorectal cancer, type 1 (120435) | Recommended to follow-up and have colonoscopy |
| 36- F | *BARD1* | Y404* | Breast cancer susceptibility (114480) | Recommended to discuss with physician and cancer genetic counselor |
| 47-M | *BRCA2* | V220fs | Breast-ovarian cancer, familial 2 (612555) | Recommended to have self- and clinical- breast exams; Discuss with cancer genetic counselor |
| 52- M | *RET* | C609Y | Medullary thyroid carcinoma (155240); Susceptibility to Hirschsprung disease 1 (142623) | Recommended to follow-up and test daughter with Hirschsprung’s disease |

\* We have (1) retained the language used by the participant and (2) included any reported family history that is plausibly related to the phenotype of concern.

Secondary genetic variation affecting cardiac genes was identified in two individuals with a previous clinical diagnosis and had a family history of cardiovascular phenotypes. One 30-year-old female reported to have experienced cardiomyopathy postpartum, had a paternal family history of arrhythmia, and her paternal uncle suffered two “heart attacks” prior to age 40. She was found to harbor a frameshift variant in *DSG2,* a gene associated with arrhythmogenic right ventricular dysplasia and dilated cardiomyopathy (MIM: 610193, MIM: 612877). Although *DSG2*has not per se been associated with peripartum cardiomyopathy (PPCM), we find it probable that the variant explains her disease history. The clinical symptoms of PPCM are similar to that of dilated cardiomyopathy ^10^ and other genetic variants associated with dilated cardiomyopathy may represent susceptibility factors for PPCM ^11^. In a 52-year-old male with hypertrophic cardiomyopathy and arrhythmia, we identified missense variation in *ANK2,* a gene associated with ankyrin-B-related cardiac arrhythmia and long QT syndrome (MIM: 600919). It is unknown whether this individual presents with long QT intervals. Additionally, although not clearly related to *ANK2* variation, this individual also reported his father had ischemic heart disease.

Finally, six of the eight parents carrying P/LP variation in *HBB* reported having sickle cell or thalassemia trait at time of enrollment (Table 2; Table S2).

### Secondary genetic variation in symptomatic individuals and/or those that have family history of disease

We identified secondary variants in 13 individuals with no previous diagnosis or genetic testing despite the manifestation of disease and/or family history (Table 3; Table S1). Given information provided at time of enrollment, six of these cases (*CLCN1, MFN2, BRCA1, BRCA2, BARD1, PMS2;* Table 3) would have met criteria to justify genetic consultation and testing via standard clinical recommendations ^12, 13^. Given additional phenotypic information acquired at return of results, two additional cases (*SCN4A, HARS;* Table 3) would have met such criteria ^14, 15^. These eight cases are described below.

A heterozygous missense variant in *CLCN1* was identified in a 29 year-old female who reported leg cramps and restless legs beginning in childhood. Variation in *CLCN1* associates with myotonia congenita (MIM: 160800) characterized by muscle stiffness and inability to relax muscles after voluntary contraction, symptoms that are exacerbated by colder temperatures. Her mother was diagnosed with myotonia congenita when she was 10 years old and her maternal grandfather had a muscle biopsy performed in his 30s due to presentation of symptoms, including “stiffness” that occurred “especially in cold [temperatures]”. In a separate case, a heterozygous missense variant in *MFN2* (Charcot-Marie-Tooth (CMT) Disease type 2A2A, MIM: 609260) was identified in a 35-year-old female who reported balance difficulties and weakness since childhood that has progressed to severe cramping, myalgia, and numbness most prominently in lower extremities, and is exacerbated by exercise. Her family history is notable for neuromuscular disorder, with similar symptoms present in her brother, father, paternal grandmother, and paternal aunt. Though a clinician has not formally evaluated her, she reported that her brother was diagnosed with CMT.

We also identified cancer risk variants in a number of individuals who report family history of cancer. We identified a frameshift variant in *BRCA1* (familial breast/ovarian cancer, MIM: 604370) in a 40-year-old male whose mother was diagnosed with breast cancer in her thirties. In another case, variation in a canonical acceptor splice site of *BRCA2* (familial breast/ovarian cancer, MIM: 612555) was identified in a 38-year-old female who had a history of breast cancer on both sides of the family - paternal grandmother (diagnosis at unknown age) and maternal grandfather (age 60). A frameshift variant in *BARD1* (MIM: 114480) was identified in a 33-year-old female whose maternal grandmother had bladder, lung, and peritoneal cancer as well as a great-grandmother diagnosed with breast cancer in her fifties. Additionally, a frameshift variant in *PMS2* (hereditary nonpolyposis colorectal cancer; MIM: 614337) was identified in a 43-year-old male with family history of colon cancer - father (diagnosed in his sixties) and paternal aunt (forties). This individual also had a paternal aunt and grandmother who were diagnosed with breast cancer in their sixties and fifties, respectively. After receipt of this finding, the study participant followed-up with a colonoscopy, found to be negative. He reports that he will continue periodic assessment.

Secondary variants were also identified in two symptomatic individuals who were not aware that their symptoms were unusual and thus never had clinical or genetic evaluation (Table 3). At enrollment, neither individual reported relevant phenotypes to the variants identified. In one case, a 28-year-old female was found to harbor a pathogenic missense variant in *SCN4A,* implicated in hyperkalemic periodic paralysis and paramyotonia congenita (MIMs: 170500; 168300), neuromuscular disorders characterized by intermittent muscle weakness and/or myotonia. At results return, she reported a history of painful stiffness during exercise that began at approximately age five and that her throat “locks up” after drinking cold liquids. Additionally, she reported that her eyelids “stick” and “become heavy” throughout the day. She noted that her mother displays similar phenotypes. This individual has plans to follow-up with a neurologist. In a second case, a 41-year-old male was found to harbor pathogenic variation in *HARS,* associated with Charcot-Marie-Tooth disease (MIM: 616625) characterized by gait difficulties and sensory impairment caused by peripheral neuropathy. At return of results this individual indicated that he was “clumsy”, discharged from military boot camp due to his inability to march in formation, and often wears out shoes because of feet shuffling.

### Identification of genetic risk factors in individuals who are currently asymptomatic and have no family history of disease

We also identified P/LP variants in individuals that are currently asymptomatic and no family history of disease (Table 3). Two unrelated individuals, a 52-year-old female and a 50-year-old male, were found to harbor variation in *SCN5A* (Long QT syndrome, MIM: 603830) and *DSG2* (dilated cardiomyopathy, MIM: 618277), respectively. A 31-year-old male study participant was found to harbor a missense variant in *ACTN1,* associated with a bleeding disorder (MIM: 615193). Finally, P/LP cancer-associated variants were identified in four participants with no personal or family history, including one in each of *MSH2, BARD1, BRCA2,* and *RET* (Table 3; Table S1). Notably, a pathogenic missense variant (C609Y) in *RET,* associated with multiple endocrine neoplasia type 2A (MEN; MIM: 171400), medullary thyroid carcinoma (MTC; MIM: 155240), and/or Hirschsprung’s disease (MIM: 142623), was identified in a 52-year-old male participant who reported no history of *RET*-associated cancer. C609Y has been observed in many MTC-affected individuals and has been designated as level B risk from the American Thyroid Association (level D is highest risk), with expected age of onset of less than 30 years ^16, 17^. Recommendations for C609Y carriers vary but often include prophylactic thyroidectomy at a young age ^18,19^. However, more recent studies indicate *RET* C609Y may have lower penetrance or later onset of MTC than previously noted ^20,21^, which are consistent with the observation of no personal or family history of cancer in this family. Interestingly, while C609Y was not transmitted to the enrolled, developmentally delayed proband, the family reported that they have another daughter (not enrolled) who has Hirschsprung’s disease and is therefore likely to have inherited C609Y. The family was referred for genetic counseling to test for the variant in the Hirschsprung’s-affected daughter and that both the father and daughter follow up with oncologists.

### Secondary findings in DD/ID-affected children

For three enrolled children, we identified secondary variation not inherited from a parent. Two individuals whose biological parents were not available harbored pathogenic variation in *CFTR* (Phe508del) and *BRCA2* (Leu579*), respectively. Also, a six-year-old female harbored a pathogenic *de novo* variant in *FBN1* (Asn2144Ser). At time of analysis, this proband did not exhibit Marfan phenotypes (MIM: 154700), with exception of crowded teeth and scoliosis. In three additional probands, compound heterozygous variation associated with recessive disease was identified. Two P/LP variants, one inherited from each parent, in *OCA2* (oculocutaneous albinism type II, MIM: 203200) were identified in an eleven-year-old male and his six-year-old brother; both presented with albinism. In a third case, a nine-year-old female with cataracts was found to inherit a P/LP variant from each parent in *FYCO1,* a gene associated with cataract 18 (MIM: 610019).

## DISCUSSION

The ACMG estimated that secondary findings in genes relevant to a defined list of actionable phenotypes (e.g. cardiac arrhythmias, cancers) would be found in ˜1% of sequenced individuals ^1,2^. We observed variation in ACMG genes in 1.4% of parent participants, consistent with that estimate and the 1%-5.6% reported by other research and clinical laboratories ^22-25^.

Our study assessed carrier status in all participants for only three genes, *HBB, HEXA,* and *CFTR,* leading to the identification of P/LP variation in ˜6.1% of parent participants. These genes were selected based on their anticipated frequencies in the population sampled and our desire to balance yield with analytical and cost burden. Had we assessed all genes known to associate with recessive disease ^26^, the burden of analysis would have increased substantially ^27,28^. Further, expanded carrier screening and discovery efforts would have increased Sanger validation costs and the time required from genetic counselors and medical geneticists for return of results. Thus, while our choice of genes as targets for carrier analysis was semi-arbitrary, the restriction to only three genes imposed minimal analytical burden and led to a substantial but manageable yield.

One additional more comprehensive carrier status strategy we used was to search within both parents of a parental pair for P/LP variants in the same gene (expanding beyond *CFTR, HBB* and *HEXA*). Of the 365 parental pairs enrolled, recessive disease risk (i.e., 25% for future children) was identified in one. This small number was likely due to our relatively stringent criteria for classifying variants of this type as pathogenic or likely pathogenic, but these numbers are likely to grow in the future as evidence accrues on the pathogenicity of variants in genes causing Mendelian disorders ^22^. The treatment of parental pairs as units of analysis for carrier status is an effective way to minimize analytical and cost burden and yet effectively capture those carrier results likely to have the greatest potential impact.

Copy-number variation (CNV) was not explored in parents as a source of secondary findings. This decision was driven by the considerable manual scrutiny that is required to evaluate the technical quality of CNVs, the costs and challenges of CNV validation, and the absence of robust CNV population frequency data, particularly for smaller events. Analyses of secondary P/LP CNVs may be of interest to future efforts to increase the overall yield of medically relevant variation within sequencing data.

### Patient preferences

The question of whether patients and research participants need to be offered choices about receiving secondary findings has been debated, especially after the release of ACMG’s original secondary findings recommendations in 2013 ^1^. Multiple studies have documented that most participants want most, and usually all, possible secondary findings. This trend is consistent between studies asking this question as a hypothetical ^29-33^ or to inform actual return of results ^34-37^. Consistent with these previous studies, the vast majority (84.8%) of parents participating in our study chose to receive all 13 categories of potential secondary results. However, a minor but substantial fraction of participants (15.2%) declined at least one category of secondary results; 1.6% declined all such results. One of the secondary findings listed in Table 3 was not returned because the parent had declined the relevant category.

### Challenges associated with variant interpretation

One of the most challenging tasks when analyzing secondary findings is interpretation of genetic variation, particularly for variants that have not been previously described in scientific literature or in clinical genetic databases. Even those variants labeled as pathogenic in variant databases are often supported by only weak underlying evidence or are even associated with strong evidence for being benign ^38^. Interpretation is made even more challenging when an individual harbors potential disease-associated variation but does not present with the associated phenotype or have a family history of disease. That said, in this study, ACMG evidence codes were assigned and variants that were deemed to be P/LP were offered for return regardless of the presence or absence of any particular phenotype or family history. Even for those with indications of disease, the particular phenotypes reported (Table 3) are not necessarily directly related to the presence of the given variant. Imprecision and incompleteness of self-reported diseases and family histories and limitations to knowledge of penetrance and expressivity for any given gene, and especially any given variant, can all make interpretation more challenging.

### Utility of secondary findings

The secondary genetic findings that we identified may be of considerable utility to the parent participants. For five individuals, we were able to confirm, and genetically explain, a previous clinical diagnosis (Table 3). Such information may prove useful for future clinical management and in discussions with family members that may carry the same variant. Secondary genetic findings were also identified in 13 individuals who reported family history or symptoms that are likely to associated with the detected variant. As described in detail in the results section, it is clear that genetic counseling and testing could/should have been performed on eight cases based on presentation of symptoms and/or family history. Additionally, we identified secondary genetic variants in four individuals who have an increased risk of disease with modest but nontrivial evidence for disease (two cases of *KCNQ1; one case each of MYBPC3* and *DDX41*). Through participation in our study, these individuals now have a better understanding of their cause or risk of disease and are in position to better manage that disease or risk of disease.

We also identified secondary genetic variation in seven individuals who report neither symptoms nor family history of disease (*MSH2, RET, BARD1, BRCA2, ACTN1, SCN5A, DSG2*). These study participants appear to be at increased risk of disease and it has been suggested that they to follow-up with an appropriate specialist (Table 3) in the hopes that actions can be taken to screen for, prevent, or mitigate unobserved disease in these individuals.

Finally, we also identified secondary variation in DD/ID affected probands that were not identified in parents, either due to unavailability of parents, (n=2) or as a result of the variant arising *de novo* (n=1). Further, three children exhibited recessive disease unrelated to DD/ID and were found to harbor compound heterozygous variation that explained their disease (i.e. albinism and cataracts).

### Challenges of returning unexpected variants to families

Many parents in this study have experienced a diagnostic odyssey in hopes of identifying the cause of their child’s developmental disabilities. Individuals who carried P/LP secondary variants therefore required counseling and recommendations for clinical follow-up regarding their secondary findings, in addition to information regarding the care and well-being of their affected children. Returning genetic information relevant to a new or unexpected disease risk may be particularly problematic when no results are found relevant to the primary indication for testing. In our study, 51% of the secondary findings identified in the parents were transmitted to the DD/ID-affected proband, and 56% of the 71 parents that harbored a secondary finding did not receive a primary result for their enrolled DD/ID-affected child. The lack of a primary result may increase the shock value of a secondary finding. A parent may expect the conversation to revolve around their child’s health but instead spends time discussing the meaning of their own disease risk and/or an additional disease risk relevant to their already affected child. This fact highlights the potential financial, emotional, and clinical implications of secondary findings that should be clearly addressed in the informed consent discussion prior to sequencing so that families are aware of all the possible outcomes of this type of testing.

## Conclusions

Our study describes the identification and return of secondary variation to parents who were subject to genomic sequencing in hopes of receiving a genetic diagnosis for a developmentally delayed child. Although the return of secondary genetic variation has been debated ^39,40^, a large majority of parent participants in this study opted to receive all identified secondary findings, regardless of disease category, suggesting that participants are generally open to receiving genetic information that may be relevant to their health. This study demonstrates the utility of returning secondary variants, as it may facilitate preventative screening for individuals who are genetically predisposed to serious diseases. This information can also be useful to individuals who have been clinically diagnosed with a specific condition but for whom the causal genetic variant has been unknown. We have also shown that secondary genetic information may lead to clinical diagnosis in individuals who have experienced symptoms related to a disorder not previously diagnosed. Some individuals also described significant family history that would have justified, but did not lead to, genetic evaluation independent of their participation in this study. Finally, our study describes a framework for identifying secondary genetic variation in a broad yet manageable manner, including a limited but productive carrier screen on only a few common recessive diseases along with a more comprehensive screen that treats parents as mate pairs. The methods and results related to secondary variation identification may be of use to other research and clinical laboratories that are conducting genomic sequencing.

## Acknowledgements

We are grateful to the families who contributed to this study. We thank the staff at North Alabama Children’s Specialists (Children’s of Alabama in Huntsville, AL), the HudsonAlpha Software Development and Informatics team, the Genomic Services Laboratory at HudsonAlpha, and the HudsonAlpha Clinical Services Lab, LLC. who contributed to data acquisition and analysis.

